# Plant Chimeras: the good, the bad, and the ‘Bizzaria’

**DOI:** 10.1101/060715

**Authors:** Margaret H. Frank, Daniel H. Chitwood

## Abstract

Chimeras – organisms that are composed of cells of more than one genotype – captured the human imagination long before they were formally described and used in the laboratory. These organisms owe their namesake to a fire-breathing monster from Greek mythology that has the head of a lion, the body of a goat, and the tail of a serpent. The first description of a non-fictional chimera dates back to the middle of the seventeenth century when the Florentine gardener Pietro Nati discovered an adventitious shoot growing from the graft junction between sour orange (*Citrus aurantium*) and citron (*C. medica*). This perplexing chimera that grows with sectors phenotypically resembling each of the citrus progenitors inspired discussion and wonder from the scientific community and was fittingly named the ‘Bizzaria’. Initially, the ‘Bizzaria’ was believed to be an asexual hybrid that formed from a cellular fusion between the grafted parents; however, in-depth cellular analyses carried out centuries later demonstrated that the ‘Bizzaria’, along with other chimeras, owe their unique sectored appearance to a conglomeration of cells from the two donors. Since this pivotal discovery at the turn of the twentieth century, chimeras have served both as tools and as unique biological phenomena that have contributed to our understanding of plant development at the cellular, tissue, and organismal level. Rapid advancements in genome sequencing technologies have enabled the establishment of new model species with novel morphological and developmental features that enable the generation of chimeric organisms. In this review, we show that genetic mosaic and chimera studies provide a technologically simple way to delve into the organismal, genetic, and genomic inner workings underlying the development of diverse model organisms. Moreover, we discuss the unique opportunity that chimeras present to explore universal principles governing intercellular communication and the coordination of organismal biology in a heterogenomic landscape.

## Introduction

Chimeric and mosaic analyses were employed long before the advent of molecular genetics to explore the fundamental principles that guide organismal growth and development. The rapid release of new model organisms that have yet to be characterized at their most basic level and the advancement of molecular techniques that enable the dissection of heterogenomic interactions begs for a reemergence of these classic tools and a re-visitation to explore the molecular coordination that underlies plant development using chimeric and mosaic approaches. We start this review by defining heterogenomicity; we then review the historical context of the chimera concept, describe experimental approaches for obtaining chimeras and mosaics, and finally discuss the future of how chimeric and mosaic studies can transform our view of organismal biology.

## Definitions

The term “heterogenomic” refers to organisms that contain heterogeneous genomes. Heterogenomic can be used to describe hybrids and allopolyploids in which independent genomes are housed within a single nucleus, as well as chimeras, genetic mosaics, and the heterokaryotic condition, in which heterogeneous genomes are housed in separate nuclei. Here, we focus on the latter case of heterogenomicity, and specifically discuss two types of heterogenomic organisms: (1) chimeras, which are formed from a conglomeration cells that originated from separate zygotes, and (2) genetic mosaics, which initiate from a single zygote and are subsequently induced or mutated into a heterogenomic state (Rossant and Spence, 1998). Grafted plants, which are formed through the physical joining of separate plant parts, are another class of heterogenomic organism that we do not discuss in this review due to the recent publication of several other reviews on this topic (we refer the reader to: Mudge et al., 2009; Goldschmidt, 2014; Albacete et al., 2015; Melnyk and Myerowitz, 2015; and Warschefsky et al., 2016).

## A brief history

Chimeras have intrigued and perplexed the scientific community for centuries. Originally recognized as“sports” (phenotypically distinct branches that arise during vegetative propagation), descriptions of chimeras first appeared in the horticultural literature in 1674, when the Florentine gardener Pietro Nati discovered the ‘Bizzaria’ growing from the graft junction of *Citrus aurantium and C. medica* (**Fig 1A-B**; Tilney-Bassett, 1991; Pietro Nati, 1674). The repeated emergence of unusual sports growing out of graft-junctions from various species combinations piqued the interest of the scientific community and generated extensive speculation about the nature of genetic inheritance and plant hybridization. In his book on “The Variation of Plants and Animals Under Domestication” Darwin proposed the theory of “graft-hybrids” wherein rootstock and scion donors can fuse at the graft-junction site to asexually generate a new hybrid (**Fig 1C**; Darwin, 1868).

**Fig 1.**
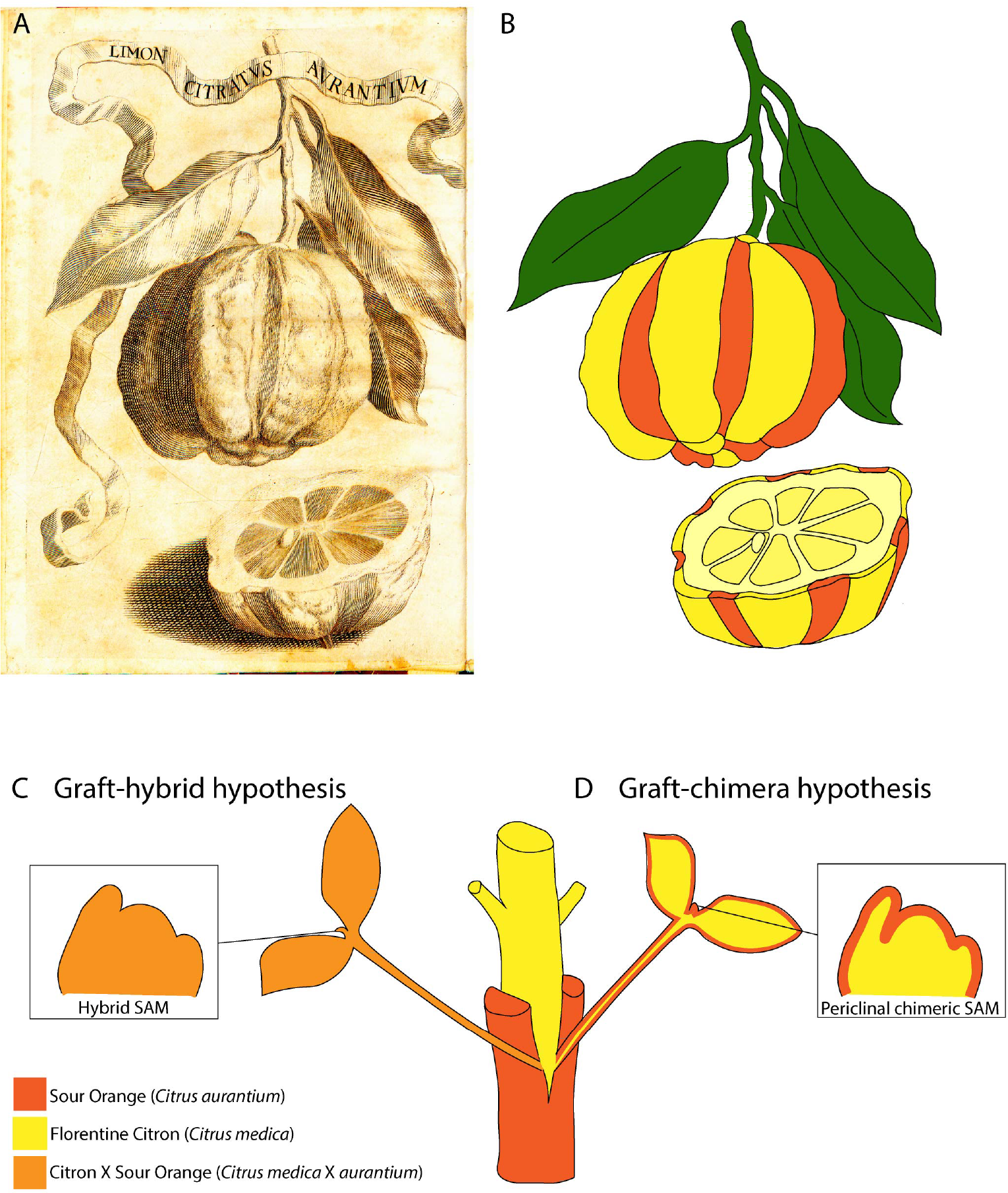
Original depiction (**A**) and genetic basis (**B-D**) of the first described chimera, The ‘Bizzaria’. Pietro Nati discovered The ‘Bizzaria’ growing as an adventitious shoot from the failed graft junction between Florentine Citron and Sour Orange. Two hypotheses, the graft-hybrid (**C**) and the graft-chimera hypothesis (**D**), were put forth to explain this unusual sport. Centuries later, Winkler (1907) and Baur (1909) demonstrated that The ‘Bizzaria’ along with many other horticultural sports resulted from a heterogeneous conglomeration (**D**) rather than asexual fusion (**C**) of parental cells.

The graft-hybrid theory was later refuted and replaced by the graft-chimera hypothesis following two foundational observations at the beginning of the twentieth century (**Fig 1D**; Tilney-Bassett, 1991). First, Winkler (1907) experimentally generated sports at the graft junction between two nightshade species, *Solanum nigrum* and *S. lycopersicum* in an effort to recapitulate Nati’s discovery and dissect the organismal basis of grafting-induced sports. While the majority of the buds resembled one parent or the other, there was one exceptional sport that grew as a longitudinal transect of the two parents, clearly indicating it originated as a conglomeration, rather than a fusion, of cells from the two species. Inspired by the fire breathing Greek monster composed of a lion’s head, a goat’s body, and a serpent’s tail, Winkler adopted the term “Chimera” to describe his morphological anomaly. Winkler (1909) also isolated several shoots that grew as phenotypic intermediates between the two parents, which he assumed were the products of cellular fusion between the progenitor species, and thus proposed that both graft-hybrids and graft-chimeras can be generated at the rootstock-scion junction. Congruent with Winkler’s work, Baur (1909) performed a series of independent experiments tracking chlorophyll inheritance in variegated geraniums, which lead him to propose a model in which mature tissues within the shoot can be traced back to clonally distinct layers from the shoot apical meristem (SAM). Following Baur’s hypothesis, virtually every sport, including Winkler’s phenotypic intermediates, could be described as a heterogeneous arrangement of cells within the SAM, negating further, serious consideration of the graft-hybrid concept.

A century later, the graft-hybrid hypothesis was re-invigorated; experiments tracking the cellular dynamics of fluorescently marked rootstock-scion combinations demonstrated that cellular and nuclear fusion does occur in rare instances at the graft junction, and can actually serve as a route for the asexual generation of allopolyploids (Stegemann and Bock, 2009; Stegemann et al., 2012; Thyssen et al., 2012; Fuentes et al., 2014). While the vast majority of grafting-induced sports are chimeras, this re-emergence of the graft-hybrid concept is a testament to the transformational power that modern techniques can have when they are applied to classical questions. It is in this light that we open our review, reopening an old topic for modern dissection.

## Classification of Chimeras and Genetic Mosaics

Chimeras and genetic mosaics can be classified by the arrangement of their genetically distinct cell types as well as the nature of their origin. Markers that allow for genotypically distinct cells to be distinguished from one another have made chimeras and genetic mosaics extraordinarily useful for performing cell lineage analyses, in which an individual cell and its descendants be tracked from a surrounding population of unmarked cells, and for teasing apart cell autonomous gene functions (in which a cellular trait is influenced by the genotype of that cell) from non-cell autonomous gene functions (in which a cellular trait is influenced by the genotype of other cells). Originally, these markers consisted of differences in the presence or absence of pigmentation (e.g. – anthocyanin or chlorophyll) or less commonly, cytological features such as genome ploidy or chromosomal rearrangements (Brumfield, 1943). Due to these limitations, the vast majority of cell lineage studies were initially restricted to chlorophyll-rich shoot systems.

Before delving into chimera classification it is necessary to give a brief overview of the structure of the shoot apical meristem (SAM). The architecture of the SAM has changed during the evolution of the land plants (Steeves and Sussex, 1989). The SAMs of bryophytes (liverworts, mosses, and hornworts) and seedless vascular plants (ferns and lycophytes) typically contain a single, conspicuous initial cell or in certain lineages (Lycopodium and Isoëtes), plural, inconspicuous initials. Most seed plants have tunica-corpus SAMs that are organized into clonally distinct cell layers with outer “tunica” layers dividing anticlinally and an inner “corpus” layer that divides both anticlinally and periclinally (Schmidt, 1924; Satina et al., 1940). Gymnosperms typically have a single tunica layer, while most angiosperms have two layers (Popham, 1951). These clonally distinct cell layers contribute to separate tissues within the newly formed lateral organs that are produced along the flanks of the SAM. In leaves of most, but not all angiosperms, the outer meristem layer (L1) forms the colorless epidermal cover, the second meristem layer (L2) forms the sub-epidermal palisade mesophyll and abaxial spongy mesophyll tissue, and the inner corpus (L3) forms the deep mesophyll and vascular tissue (reviewed in Tilney-Bassett, 1991). Thus, plants that are composed of heterogeneous cells in the SAM can be categorized based on the genetic composition of their shoot meristem layers: (1) periclinal chimeras that have a uniform, genetically distinct layer of cells in the shoot apical meristem (SAM) (**Fig 2A**), (2) mericlinal chimeras have a heterogenomic population of cells within a single layer of the SAM (**Fig 2B**), and sectorial chimeras that either have a heterogenomic population of cells traversing multiple SAM layers (**Fig 2C**) or have non-patterned heterogenomic patches of cells (**Fig 2D**). There is a vast array of techniques that are available for producing periclinal, mericlinal, and sectorial chimeras. Some methods are extremely accessible and have been employed for centuries, while others involve advanced transgenic techniques (**Table 1**). The remainder of this review not only highlights methods for creating chimeras and genetic mosaics, but also discusses the biology of these unique organisms, how they have shaped modern plant development, and their potential to transform future applications when combined with new technologies.

**Fig 2.**
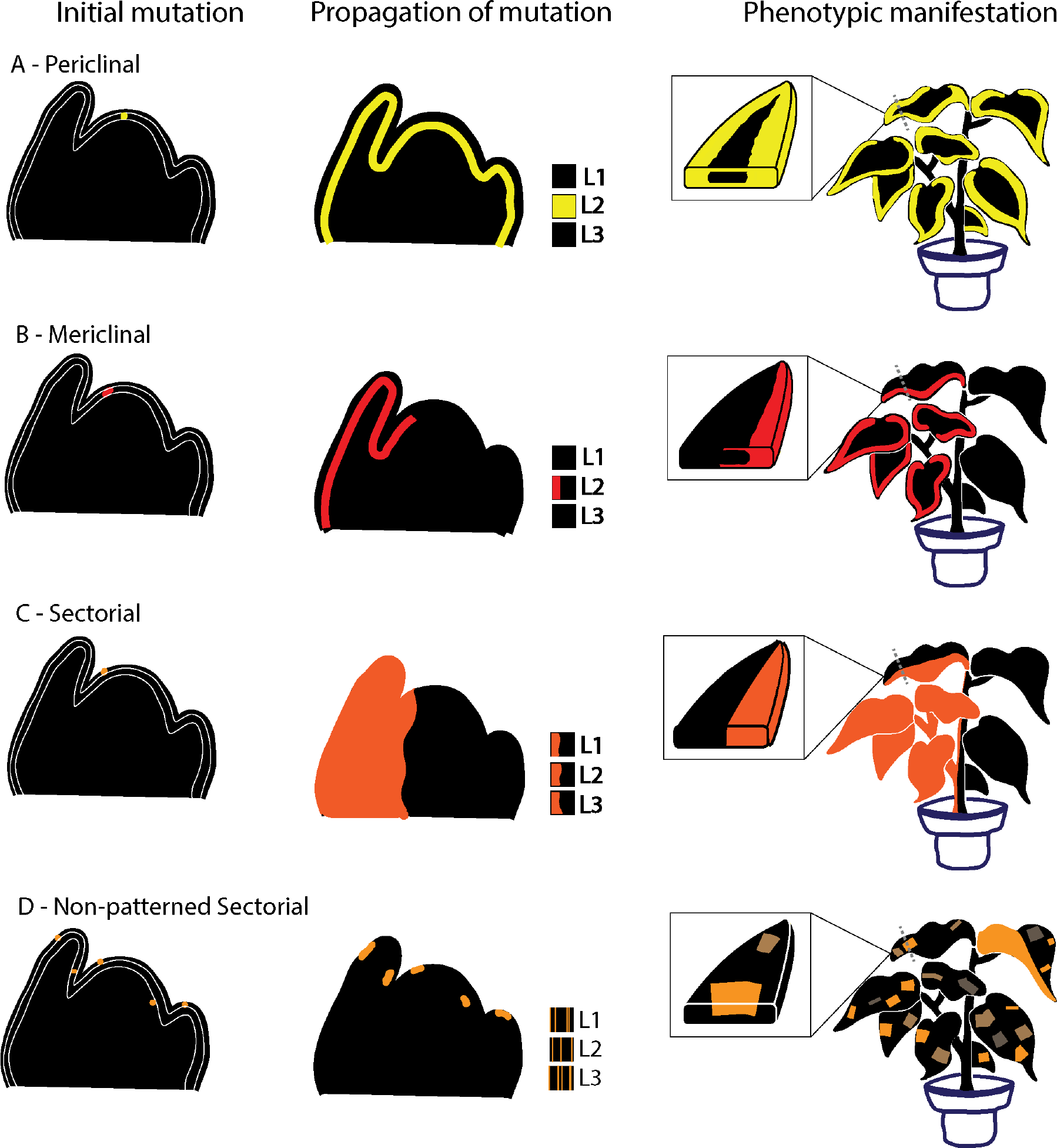
Mutational basis and phenotypic manifestation of periclinal (**A**), mericlinal (**B**), and sectorial (**C-D**) mosaics. Periclinal (**A**) and mericlinal mosaics can arise through mutations in shoot apical meristem (SAM) initial cells (shown in yellow and red, respectively). Periclinal mosaics are formed when the mutation propagates throughout the meristem layer, creating a uniform, genetically distinct stratum of cells (**A**). Mericlinal mosaics, on the other hand, arise from incomplete invasion of the meristem initials, creating a genetically sectored shoot (**B**). In this example, an L2 mutation in yellow (**A**) and/or red (**B**) gives rise to mutant features along the leaf margins and wild-type features within the leaf core. Genomically unstable plants, such as individuals with active transposons, can also give rise to sectored mosaics (**C-D**). These mosaics are often characterized as being unstable and can take the form of large sectors that traverse all layers of the shoot meristem (**C**) or have a non-patterned variegated appearance (**D**). The size and frequency of sectoring is a function of transposon (or other mutagen) activity and the rate of cell division.

**Table 1.**
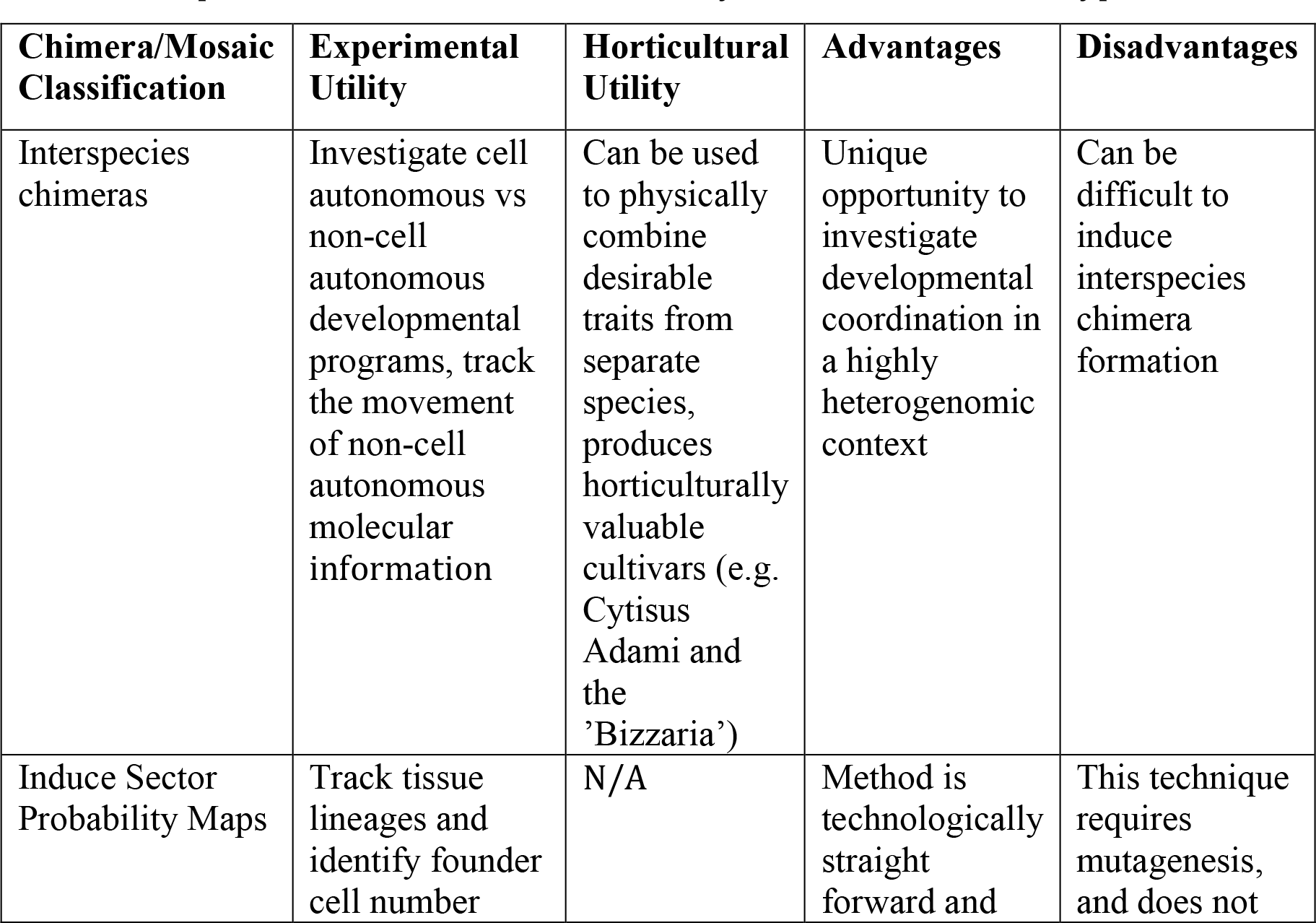
Experimental and horticultural utility of different chimeric types

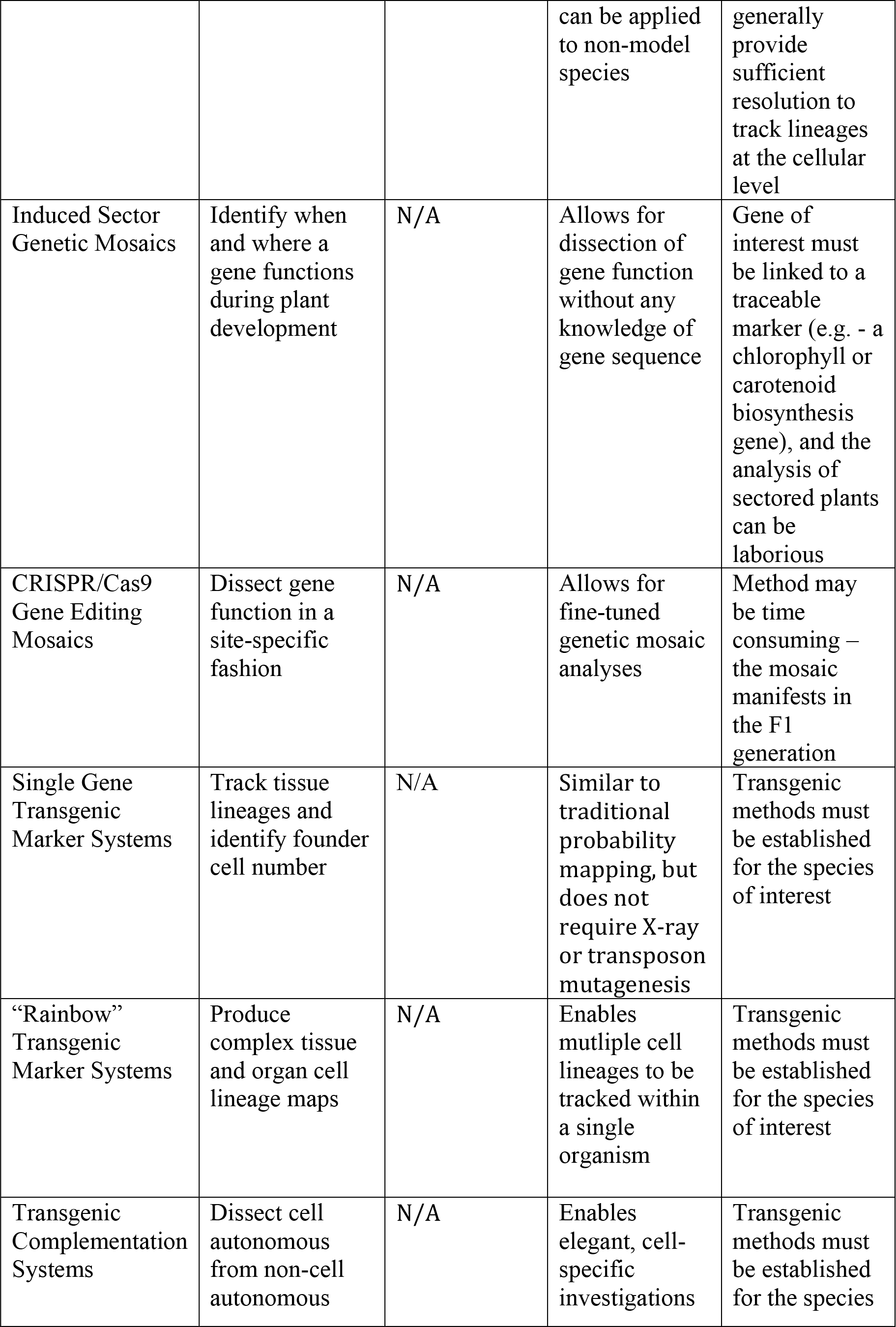

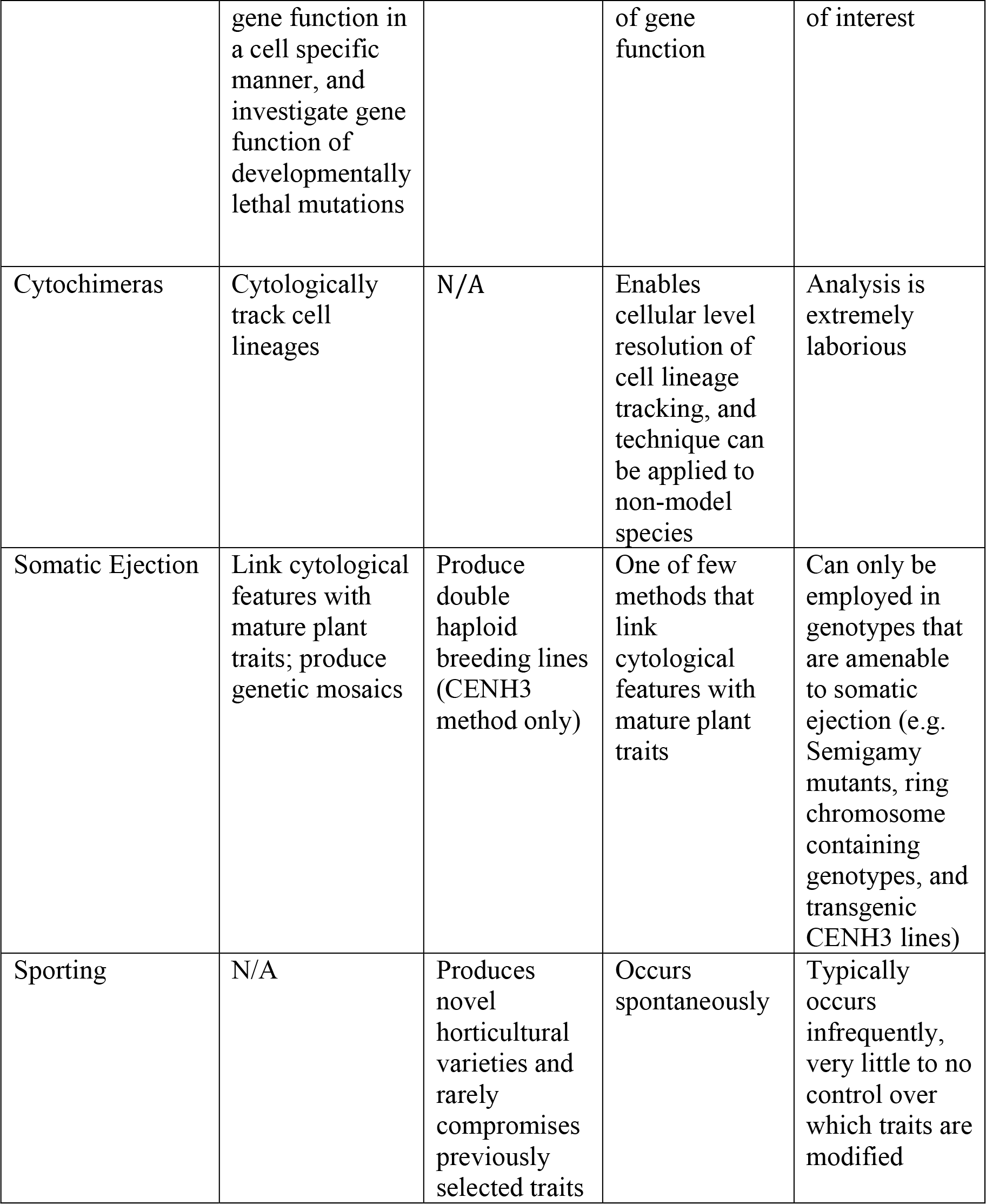

## Interspecies chimeras

The fact that fully-functioning, reproductive organisms can form out of a conglomeration of cells from species that have diverged over millions of years is a testament to the remarkable flexibility of plant development. These fantastic interspecies chimeras can be used to determine which cell layers control overall plant architecture (Kaddoura and Mantell, 1991), distinguish cell autonomous from non-cell autonomous developmental programs (Zhou et al., 2002), screen for the movement of molecular information between cell layers, and challenge the coordination of growth and development between divergent species within a single organism (Jørgensen and Crane, 1927; Marcotrigiano, 2010). Interspecies chimeras are generated by mixing cells from separate species together into a single callus culture. Heterogenomic callus is commonly formed by grafting two species together and subsequently inducing bud formation at the graft-junction (generating a graft-chimera) (Winkler, 1907; Chen et al., 2006) or by co-culturing the two species, using a modified tissue culture protocol (Murashige and Skoog, 1962; Marcotrigiano, 1984). A small percentage of the shoot meristems that emerge from these calli are composed of cells from both species. Both mericlinal and periclinal chimeras can arise from this technique and the nature of the chimera depends on the organization of the SAM. This method was inadvertently used to produce the ‘Bizzaria’, and was later employed by Winkler (1907) as a tool to provide the first experimental evidence for the cellular basis of the chimera concept (Tilney-Bassett, 1991).

The generation of interspecies chimeras is a largely underappreciated, yet powerful tool that can be used to tease apart intercellular coordination during development. For example, Zhou et al. (2002) were able to distinguish cell autonomous from noncell autonomously specified morphological, metabolic, and size features using periclinal chimeras between *Citrus sinensis* and *C. natsudaidai*. Their work revealed that epicarp and juice sac coloration is a cell autonomous trait, whereas epidermal patterning, metabolic features, and organ size are all products of inter-tissue communication (Zhou et al., 2002). The generation of periclinal chimeras between the simple-and complex-leaved species *Solanum luteum* (S) and *S. lycopersicum* (C) (respectively) played an important role in demonstrating tissue-layer coordination during leaf development (reviewed in Szymkowiak and Sussex, 1996). Leaves formed from the CSS periclinal arrangement developed simple leaves, whereas those formed from the SCC arrangement produced complex leaves (Jørgensen and Crane, 1927), indicating that the sub-epidermal cell layers largely control leaf complexity. Intriguingly, the complex leaves formed from the SCC chimeric organization developed fewer leaflets than uniform CCC plants, implying that while the L1 does not determine simple versus complex leaf formation, it can modulate the degree of complexity (Jørgensen and Crane, 1927). More recently, Marcotrigiano (2010) tested the influence of cell division and expansion on overall organ size by generating periclinal and mericlinal chimeras between the big-and small-leaved species, *Nicotiana tabacum* (B) and *N. glauca* (S) (respectively). Leaf lamina formed from the BSS genotype were big in size, whereas lamina formed from the SBB genotype were small in size, indicating that the epidermal cell layer largely controls overall organ size in leaves. This study also demonstrated that epidermal genotype can influence cell division rates in the mesophyll layer - more cells were formed in the mesophyll cell layer of BSS than SSS plants at developmentally equivalent time points. Moreover, the study showed that cell-to-cell regulation of organ size and mesophyll division rate occurs between but not within cell layers - as leaves that were genotypically split along the margin (SBB/BBB or BSS/SSS) lacked coordination for organ size and cell number between the two halves of the leaf.

## Beneficial heterospecies combinations

The utility of interspecies mosaics extends to the field, where periclinal chimeras between wild and domesticated genotypes serve as a means to physically combine beneficial adaptations from wild species with the edible or otherwise useful products of domestication. One particularly successful implementation of this strategy is in cassava, where the combination of the wild species *Manihot fortalezensis* with the cultivated species *M. esculenta* has been shown to boost edible root size by a factor of seven-fold (Nassar and Bomfim, 2013). A similar approach to plant improvement has been demonstrated for tomato where the epidermis of the wild tomato species *S. pennellii* is sufficient to confer aphid resistance to a domesticated inner core (Goffreda et al., 1990).

Once restricted to phenotypic observations, the molecular signaling that underlies the coordinated growth and development between interspecies tissue layers can now be examined with high-throughput sequencing techniques. For example, the *S. pennellii* (L1) - *S. lycopersicum* (L2/L3) chimera that confers aphid resistance was later used as a tool to profile for cell layer-specific gene expression in the shoot apex (Filippis et al., 2013). Taking advantage of single nucleotide polymorphisms (SNPs) that differentiate *S. lycopersicum* from *S. pennellii* coding sequences, Filippis et al. (2013) used bioinformatic tools to construct a map of L1-derived versus L2/L3-derived transcripts from RNA-sequence profiles of whole SAMs. This study represents one of the many ways in which clonally marked cells can be used to extract cell type-specific molecular information without having to physically sample at the sub-organ level. This method assumes that transcripts generally behave in a cell-autonomous manner; however, recent work indicates that long-distance RNA movement may be pervasive in the plant world (Thieme et al., 2015; Kim et al., 2014). While there are specific cases of cell-to-cell RNA movement (Benkovics and Timmermans, 2014; Chitwood et al., 2009; Carlsbecker et al., 2010; Knauer et al., 2013), the extent to which these “mobile molecular maps” operate on the intertissue (i.e. tissue-level) scale and whether or not mobile transcripts contain a “zip code” that specifies targeted delivery (Haywood et al., 2005), or are passively transported as a function of transcript abundance (Calderwood et al., 2016) remains poorly understood. One promising tool for teasing apart intercellular molecular movement would be to generate interspecies chimeras between fluorescently labeled individuals and subsequently screen for the exchange of molecular information between tissue layers using fluorescent associated cell sorting (FACS) in combination with transcriptomic and proteomic profiling. Such an approach would reveal, on a genome-wide level, the mass exchange of molecular information underlying the coordination of plant development, which is typically hidden in nonchimeric plants.

## Probability maps

Induced-sectoring (clonal analyses) with ionizing radiation (such as X-rays, gamma-rays, or fast neutron sources) or active transposon lines (such as Mutator and Activator-Dissociator systems) (Becraft, 2013) can be used to generate all three classes of genetic mosaics (**Fig 2**), and provides a technologically simple method for cell lineage and genetic mosaic analyses. This technique relies on the disruption of a marker gene within an otherwise homogeneous population of cells, producing marked cell lineages that can be visually distinguished from un-mutated cells, and tracked through developmental time. Genes in pigmentation pathways are typically targeted for induced-sectoring; however, radiation-induced cytological sectors have also played an important role in distinguishing cell populations in un-pigmented organs such as roots (Brumfield, 1943). The power to track a cell and its descendants over developmental time has been instrumental in shaping our fundamental understanding of the processes that guide both plant and animal development.

In contrast to animal cells where cell fate is acquired relatively early in development, plant cells tend to be highly plastic and capable of changing their developmental trajectory during late stages of differentiation (Hernandez et al., 1999; Szymkowiak and Sussex, 1996; Poethig, 1997; 1989; Irish, 1991; Dawe and Freeling, 1991). This paradigm is supported by observations of cellular invasions, in which marked cells from one tissue layer will invade and subsequently adopt the cellular identity of adjacent cells (Dermen, 1948; 1949). For example, a cell from the L1 layer of the SAM may invade the L2; this migrant along with its descendants will mature into mesophyll rather than epidermal tissue. The labile-nature of plant cell fate lead Irish and Sussex (1992) to suggest using the term “probability” rather than “fate” mapping, as position rather than lineage plays a larger role in determining the identity of a plant cell. The difference between “fate” versus “probability” mapping marks a conceptual rather than an experimental separation between plant and animal cell lineage analyses; the use of the word “probability” acknowledges that even late in their development, plant cells may take on a different cell fate depending on the context of their cellular neighborhood.

Cell lineage mapping has provided the foundation upon which all further research into the dynamics of organogenesis during plant development has been built. Cell-lineage studies carried out during the embryonic stages of development demonstrate that the entire aerial portion of the angiosperm body is derived from just a few initials (Steffensen, 1968; McDaniel and Poethig, 1988; Jegla and Sussex, 1989; Irish and Sussex, 1992; Furner and Pumfrey, 1992; Bossinger et al., 1992; Saulsberry et al., 2002), whereas analyses at later developmental stages indicate that there is considerable variation in founder cell number during determinate organ recruitment. While hundreds of cells are recruited during leaf initiation (Poethig, 1997; Stein and Steffensen, 1959; Poethig and Sussex, 1985; Poethig and Szymkowiak, 1995; Dolan and Poethig, 1998a), as few as two initials are involved in leaflet initiation (Barkoulas et al., 2008).

Newly sequenced model species with unique morphologies have inspired a revival in the use of probability mapping. Studies in ancestral lineages of the land plant phylogenetic tree show that these anciently derived species differ dramatically from angiosperms in their developmental strategies. For instance, in contrast to the hundreds of cells that are involved in angiosperm leaf recruitment, leaf-like organogenesis in the model moss, *Physcomitrella patens*, is largely orchestrated through the asymmetric divisions of a single-celled SAM (Harrison et al., 2009). Likewise, leaves in the seedless vascular plants, *Nephrolepsis exaltata* and *Selaginella kraussiana*, are derived from just one or two cells (respectively) situated along the epidermal flanks of a multicellular SAM (Harrison et al., 2007; Sanders et al., 2011). Cell-lineage mapping has also enabled quantitative explorations of novel structures in model angiosperms. The Coen lab, for instance, has combined clonal sector analyses with computational modeling, and developmental genetics, to investigate pattern formation in morphologically complex *Antirrhinum* flowers (Vincent, et al., 1995; Rolland-Lagan et al., 2003; Green et al., 2010). An extension of this strategy to diverse model systems will play a useful role in deducing the rules that underlie the formation of novel organ morphologies.

## Genetic mosaics

Induced sector analyses have also been used to investigate mutants in a tissue-and organ-specific fashion. These studies rely on the physical linkage of a marker gene with a mutant locus of interest. Heterozygous individuals are treated to produce chromosomal breakage and/or loss, unmasking hemizygous (loci that exist in a single copy state because they lack an allelic complement on the sister chromosome) mutant and wild-type sectors that can be tracked in a cell- and tissue-specific manner by observing the presence or absence of the marker (Dawe and Freeling, 1991). Genetic mosaic analyses of classical developmental mutants from maize, tobacco, and cotton have been used to decipher when and where a gene functions without any pre-requisite knowledge of the gene sequence. Barbara McClintock (1932) was among the first to experimentally demonstrate this phenomenon when she was investigating the correlation between variegated cell lineages and the somatic elimination of ring-shaped chromosomes in maize. McClintock’s initial observation has been greatly expanded upon and refined into a tool that can be used to track mutant alleles in a cell-lineage specific fashion. In a pioneering example of this technique, Johri and Coe (1983) used X-rays to unmask mutant sectors of the classic maize inflorescence mutants - *ramosa-1, tunicate, tassel-seed*, and *vestigial* - to demonstrate that in all cases these genetic factors function in a cell autonomous fashion to control tassel differentiation late in development. Since this early study, induced sectoring has proven to be particularly effective for studying the post-embryonic effects of embryo lethal mutants (Candela et al., 2011; Fu and Scanlon, 2004; Becraft et al., 2002; Neuffer, 1995), separating cell autonomous from non-cell autonomous gene functions (Hake and Freeling, 1986; Hake and Sinha, 1993; Sinha and Hake, 1990; McDaniel and Poethig, 1988; Scanlon and Freeling, 1997; Becraft et al., 1990; Becraft and Freeling, 1991; Becraft et al., 2002; Scanlon, 2000; Furner et al., 1996; Dolan and Poethig, 1998b; Dudley and Poethig, 1993; Dudley and Poethig, 1991; Foster et al., 1999; Szymkowiak and Irish, 1999), and comparing adjacent mutant and wild-type cell types within a single developmental stage (reviewed in Neuffer, 1995). The success of induced sector analysis relies heavily on the selection of appropriate marker genes and the mutagenesis strategy (the following reviews cover in-depth discussions concerning these parameters: Hake and Sinha, 1994; Neuffer, 1995; Becraft, 2013). The optimization of these parameters can be circumvented using an inducible transgenic mosaic system (discussed below).

## Transgenic marker systems

In recent years, elegant transgenic systems have greatly extended cell lineage and genetic mosaic analyses to virtually every tissue type during the plant life cycle (Sieburth et al., 1998; Kidner et al., 2000; Sessions et al., 2000; Jenik and Irish, 2001). While there are a wide variety of engineered combinations they all generally consist of an inducible marker whose expression can be stochastically “flipped” (turned on through the excision of an interrupting sequence or turned off through excision of the marker itself) (**Fig 3A**). Typically, beta-glucoronidase (GUS) or a fluorescent protein is used as the marker and the Cre-Lox recombinase or the Activator/Dissociator (Ac/Ds) transposon system is used to flip marker gene expression. Jenik and Irish (2001) constructed an adaptation to this transgenic reporter strategy, using the Activator Dissociator (Ac/Ds) transposon system, to generate genetic mosaics of the petal and stamen homeotic mutant, *apetala3 (ap3)*. They introduced a complementation construct into an *ap3* mutant background in which a wildtype copy of *AP3* constitutively driven by 35S was bordered by Ds sequences and followed by a promoterless *GUS* reporter. Constitutively expressed Ac driven under a 35S promoter lead to the stochastic excision of the Ds bordered *AP3* gene, generating mutant sectors that were marked by the recovery of *GUS* expression. This study revealed that *AP3* functions in both the cell autonomous patterning of epidermal cells and non-cell autonomous coordination of organ shape, and demonstrated the efficacy of transgenic mosaic systems for observing cell lineages in organs that lack chlorophyll markers.

**Fig 3.**
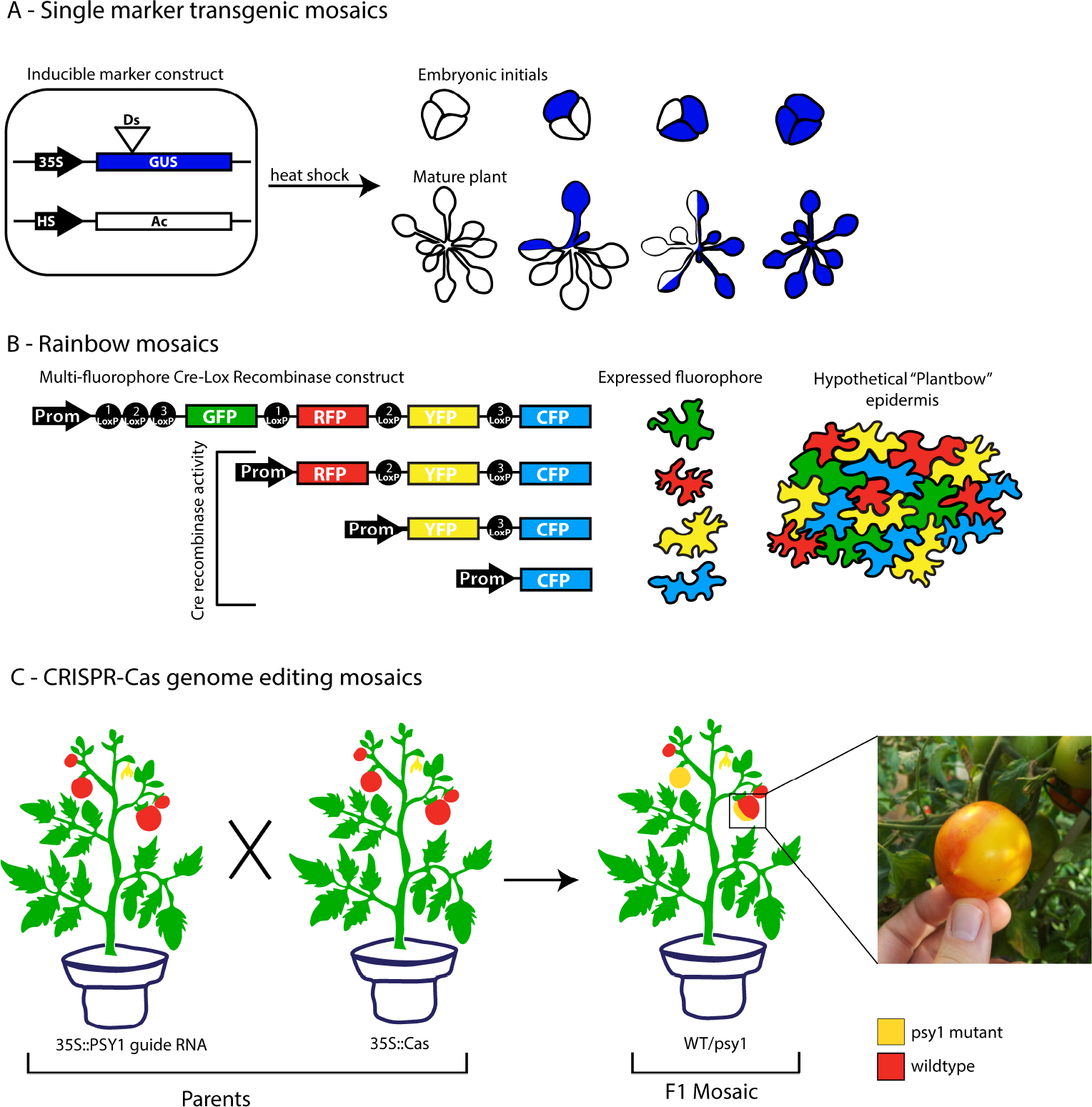
Examples of single marker,“Rainbow” and CRISPR-Cas transgenic mosaic systems. (**A**) Illustration of the single marker mosaic system used by Saulsberry et al. (2002) to map embryonic cell fate in Arabidopsis. In this example GUS expression is interrupted by the *Dissociator (Ds)* transposon, and heat shock induction of *Activator* (*Ac*) expression leads to the recovery of GUS expression through the *Ac*-promoted excision of *Ds.* The full spectrum of resulting mosaic embryonic initials as well as the mature plants that arise from each mosaic combination are illustrated from left to right with no excisions (white plant), one excision (1/3 blue), two excisions (2/3 blue), and three excisions (blue plant). (**B**) Schematic of the original Brainbow system that was used to mark developing neuorons in the mouse brainwith four independently expressed fluorophores (Weissman et al., 2011). The Brainbow system relies on Cre-recombinase excision at three different LoxP sites leading to stochastic expression outputs. In this example, excision at LoxP-1 leads to the loss of GFP and the expression of RFP, whereas excision at LoxP-2 leads to the loss of both GFP and RFP and thus the expression of YFP, and the excision of LoxP-3 leads to the loss of GFP, RFP, and YFP, leaving GFP to be expressed. Since the inception of Brainbow, several modifications have been made, including the insertion of multiple fluorophore constructs, which allows for combinatorial expression outputs and greatly extends the systems ability to unambiguously mark individual cells (e.g. – RFP plus CFP = Purple Fluorescence) (Weissman and Pan, 2015). Here, we propose a hypothetical “Plantbow” system using the plant epidermis as a model, in which individual cells within any tissue layer can be fluorescently marked and tracked for the life of the cell. (**C**) CRISPR-Cas DNA editing technology offers a new mechanism for generating site-specific genetic mosaics. Here we illustrate an example from the Levy lab (2016, personal communication) in which F1 targeted DNA editing mosaics can be generated by crossing together guide RNA and Cas9 expressing parents. In this example, the Phytoene Synthase1 (PSY1) gene, which results in yellow fruits when it is knocked out, is targeted by the guide RNA and a picture of the mosaic red (wildtype) and yellow (psy1) tomato fruit is shown.

Several powerful adaptations of transgenic mosaic methods have been designed for animal systems. G-TRACE combines a Gal4-enhancer trap with Gal4-dependent fluorescent reporters, allowing for the simultaneous and real-time examination of gene expression and cell lineage patterns within any Drosophila tissue (Evans et al., 2009). “Brainbow” is another system that relies on the stochastic flipping of multiple fluorescent proteins, creating combinatoric fluorescent outputs that can be used to distinguish the individual cells in a wide variety of animal cell systems (Weissman and Pan, 2015). Brainbow was originally invented to track individual neurons in the mouse brain (Weissman et al., 2011), and has since been extended to a wide variety of cell-type specific animal systems, including imaginal wing discs in drosophila (“Flybow” Hadjieconomou et al., 2011) and skin cells in zebra fish (“Skinbow” Chen et al., 2016). It is easy to imagine how similar tools for model plants could greatly enhance our examination of complex developmental questions, such as position-dependent cell differentiation. For example, the G-TRACE system could be used to tease apart the interplay between gene expression dynamics and the acquisition of position-dependent cell fate. Here, we propose a “Plantbow” system that would enable high resolution in *vivo* tracking of cell lineages, which could ultimately improve our understanding of how complex morphological structures are formed in new model species (**Fig 3B**).

CRISPR-Cas gene targeting, which allows for specific sequences within the genome to be edited (Bortesi and Fischer, 2015), offers a new mechanism for generating sequence-specific genetic mosaics. CRISPR technology relies on the co-expression of a guide RNA that targets the genomic region of interest and a Cas endonuclease that edits the targeted DNA. Levy et al. (2015, personal communication) discovered that crossing Cas and guide RNA expressing plants together results in F1 targeted editing mosaics (**Fig 3C**). While this advanced genetic mosaic system has yet to be widely utilized, it offers a promising approach to finely dissect gene function on the nucleotide site-specific level.

## Cytochimeras

Cytochimeras are mosaics that differ in their cytological features (typically nuclear size), can be synthesized through induced-polyploidization with colchicine treatment (Dermen, 1940). While photosynthetic markers allow for efficient lineage tracking at the organ level, cytochimeras provide a high-resolution method of tracking individual cells at a histological level. This technique is rarely used today, due to the labor-intensive methodology required to examine these mosaics; however, the knowledge garnered from these classic studies still functions as the cornerstone upon which all other investigations of the shoot apex are built. The demonstration that the SAM is organized into clonally distinct cell layers, and subsequent tracking of these layers into tissue positions within mature organs, is based upon impressive, histological dissection of cytochimeras (Satina et al., 1940; Dermen, 1947, 1953; Satina and Blakeslee, 1941, 1943; Baker, 1943; Satina, 1944; Stewart and Burk, 1970; Stewart and Dermen, 1970, 1975). Moreover, the realization that plant cells lack fixed developmental fates was unequivocally observed in periclinal cytochimeras, in which layer invasions could be tracked all the way from their origin in the SAM into mature organ tissues (Poethig, 1987; Dermen, 1953). In contrast to the extensive cytochimeric analyses that have been carried out in angiosperms, there is little to no information concerning the clonal organization of cells within seedless vascular plant meristems (Kawakami et al., 2007). The distinct apical-cell type SAM organization of species found within these early land plant lineages makes them prime candidates for cytochimeric tests of simple hypotheses concerning the functional relevance of the conspicuous apical cell (Steeves and Sussex, 1989).

## Somatic ejection

The stochastic loss or change of unstable genetic material during mitosis can also lead to the formation of chimeras. Genetic instability occurs quite frequently in plants that have unstable chromosomes (such as ring chromosomes), transgenic plants with altered CENH3 coding sequences (Ravi and Chan, 2010), chromosomes that are prone to somatic recombination (reviewed in Dawe and Freeling, 1991), or in mutants that distort nuclear dynamics during mitosis (Turcotte and Feaster, 1963). The *semigamy* mutant in cotton offers a particularly useful system for tracing cell lineages based on the random ejection of genetic material. *Semigamy* egg cells are defective in paternal-maternal nuclear fusion during fertilization. Crossing a genotype of interest onto *Semigamy* gives rise to double haploid offspring that stochastically lose either of the parental nuclei, creating genetic mosaics between the paternal and maternal genomes (Turcotte and Feaster, 1963; 1967). Dolan and Poethig (1991 and 1998a,b) leveraged this elegant genetic setup to perform classic cell lineage analyses in cotton and investigate the tissue-specific functions of the dominant, complex leaf mutant *Okra*. Their analyses showed that genetic mosaics with the *Okra* mutation in any of the three SAM layers could lead to partial expression of the *Okra* mutant phenotype, demonstrating that the compound leaf phenotype is a product of *Okra* function in all three layers of the shoot apex (Dolan and Poethig, 1998b).

## Sporting

Waiting for a chance mutation is undoubtedly the easiest way to obtain a genetic mosaic. Somatic mutation rates vary considerably at the species level; however, Lynch et al. (2010) were able to use data from Arabidopsis mutation accumulation lines (Ossowski et al., 2010) and the estimation that each generation is separated by 40 cell divisions (Hoffman et al., 2004) to calculate that approximately 1.6 × 10^−10^ mutations occur per base per cell division. According to these estimates, one mutation occurs approximately every six cell divisions; meaning that the average Arabidopsis plant is composed of a heterogenomic population of cells and is in essence a genetic mosaic long before it germinates (Ossowski et al., 2010; Lynch, 2010). The majority of these mutations occur in non-coding regions of the genome (approximately 75% of the Arabidopsis genome consists of non-coding sequence; Bevan et al., 2001), or occur in cells that are close to differentiation, thus having little to no obvious affect on plant phenotype. However, rare somatic events can create drastic organismal changes. These serendipitous mutations that occur in meristematic cells give rise to “bud-sports”, which are defined as somatic events that produce morphological shifts during vegetative development.

“Sporting” is the innovative force behind the isolation of new grape varieties (Skene and Barlass, 1983), the cultivation of thornless blackberries (Darrow, 1931; McPheeters and Skirvin, 1983), and the tremendous diversity of variegated cultivars that populate horticultural nurseries (covered in Tilney-Bassett, 1991). While most horticultural sports have yet to be traced to their molecular origins, the viticulture community has been particularly successful in identifying the mutational basis of economically important somatic events. For example, the *Vitis vinifera* variety Pinot Meunier, which happens to be one of only three grape varieties predominately used for Champagne production, arose spontaneously from a Pinot Noir shoot. Subsequent tissue culture isolation of L1 and L2 layers from Pinot Meunier revealed that this new variety was generated through a dominant negative mutation in a DELLA repressor within the L1 layer of the SAM (Boss and Thomas, 2002). Regeneration from just the L1 layer gave rise to yet another variety called “Pixie” (Cousins, 2007). Pixie is a dwarfed version of Pinot Meunier that bears inflorescences in place of tendrils and flowers earlier than its chimeric counterpart, making it an ideal new model system for carrying out genetic studies in grape (Boss and Thomas, 2002). Berry color is another highly dissected trait that sports quite frequently during grape propagation. Extensive work on the genetic and genomic underpinnings of berry color have shown that nearly all Pinot berry variants can be mapped back to a single locus consisting of four tandem MYB transcription factors on chromosome 2, aptly named the“berry color locus” (Migliaro and Crespan, 2014; Walker et al., 2006; 2007; This et al., 2007; Vezzulli et al., 2012; Azuma et al., 2011; Fournier-Level et al., 2010; Furiya et al., 2009; Kobayashi et al., 2004; Yakushiji et al., 2006). Somatic mutations within the berry color locus are frequently maintained within a single layer of the SAM; thus, berry pigmentation is an integrated output of both the allelic nature of the locus and the SAM layer in which the allele is expressed (Walker et al., 2006; Hocquigny et al., 2004; Shimazaki et al., 2011).

## Variegation in the garden

Variegated mosaics, which are defined as plants with pigmentation patterning due to SAM layer-specific mutations, provide another horticulturally abundant example of sporting. Striking leaf and flower color variants are inescapable in modern landscaping. These plants have two main genetic origins; they are either periclinal mosaics that arose through a stable meristem-layer specific mutation in a crucial component of a pigmentation biosynthesis pathway (**Fig 4A-D**) or they are nonpatterned sectorial mosaics with highly active transposons that hop in and out of pigmentation biosynthesis genes creating random sectors (**Fig 4E-G**; Tilney-Bassett, 1986). Beyond beautifying the garden, variegated plants have played an instrumental role in the establishment of the chimera concept. Baur’s (1909) investigation of leaves in variegated geraniums lead to a model in which clonally distinct layers of the SAM contribute to independent cellular regions within the differentiated leaf. In leaves, the L1 forms the unpigmented epidermal layer, the L2 forms the upper and lower mesophyll and is particularly pronounced along the margins of the leaf, and the L3 forms the core mesophyll and vascular tissue and is visible in the center of the leaf (**Fig 2A; Fig 4A**). This clear and consistent visual readout of SAM organization makes it easy to observe cellular invasions within the SAM. For example, a mosaic with a green L2 and white L3 that produces leaves with a green margin and white center (**Fig 4B**) may sport into an albino shoot if the L3 invades the L2 (**Fig 4C**), or recover into an entirely green shoot if the L2 invades the L3 (**Fig 4D**). Often times, these SAM layer transitions occur in stages where one or two meristem initials are replaced by a neighboring layer, giving rise to intermediate mericlinal mosaics in which half or a quarter of the variegated plant sports into a new pattern of variegation (**Fig 4H-J**). A century of continued exploration following Baur’s initial proposal has shown that predictable pigmentation patterns can be mapped for virtually any species and any organ-type, making these botanical puzzles incredibly useful for investigating cell lineages and SAM dynamics in non-model species.

**Fig 4.**
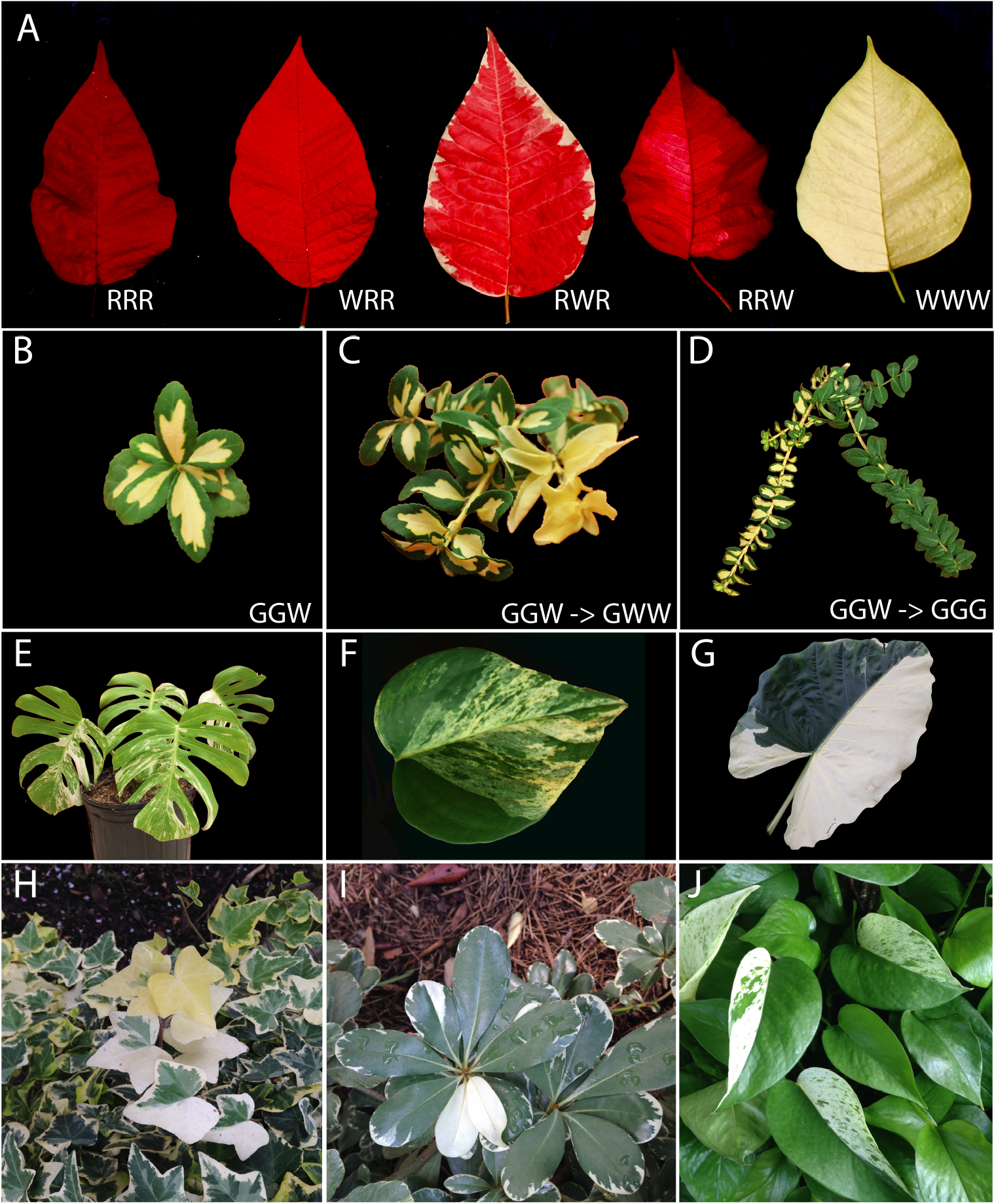
Examples of sporting in variegated mosaics. A full spectrum of potential red (R) and white (W) periclinal arrangements in the showy foliage of *Euphorbia pulcherrima* ‘Poinsettia’ is shown in (**A**). A periclinal green-L2 white-L3 *Euonymus fortunei* ‘Moon Shadow’ (**B**) sports into an albino shoot when the L3 invades the L2 (**C**) and recovers into a uniform green shoot when the L2 invades the L3 (**D**). Active transposons in non-patterned variegated varieties create frequent G/W sectoring in *Monstera deliciosa* ‘Variegata’ (**E**), *Epipremnum aureum* ‘Golden Pothos’ (**F**), and *Alocasia macrorrhizos*’ ‘Variegata’ (**G**). Sporting frequently occurs through incomplete invasion of SAM initials, giving rise to mericlinal sectors. Here, we show mericlinal invasions of *Hedera helix* ‘Gold Child’ sporting from GWG to GWW (**H**), *Pittosporum tobira*’ ‘Variegata’ GWG with a mericlinal sector of GWW (**I**), and mericlinal *Epipremnum aureum*’ ‘Marble Queen’ (**J**).

## Microchimerism

Mutations that spawn-striking shifts in plant phenotype are widely appreciated as important drivers of evolutionary change and agronomic advancement; in contrast, the collective impact of small effect somatic mutations remains poorly understood. In annuals, these events may have little to no impact on the life cycle of the plant. However, *very* long-lived organisms that collect somatic mutations for thousands of years such as the Bristle Cone Pine and the Llangernyw Yew (Sussman and Zimmer, 2014), or long-lived clonal lineages such as the creosote bush and box huckleberry, present intriguing systems for investigating the beneficial versus detrimental tradeoffs of somatic events. Evidence for the beneficial impact of somatic mutations comes from evolutionary modeling of plant-herbivore interactions between long-lived trees and rapid-cycling herbivores, which indicates that somatic mutations provide a viable mechanism for slow-cycling plants to evolve and defend against their attackers (Folse and Roughgarden, 2012; Whitham and Slobodchikoff, 1981). Next-generation sequencing technologies that allow for deep genomic profiling of isolated cell populations have proven to be hugely successful in advancing our understanding of somatic mosaicism and its relation to human diseases (Pagnamenta et al., 2012; Yamaguchi et al., 2015). Now, a similar approach has been applied to long-lived plants, providing experimental support for the theoretical model of uni-generational evolution. Transcriptomic profiling of mosaic sectors within Eucalyptus individuals demonstrates that herbivore resistant patches are marked by massive transcriptomic remodeling that is associated with enhanced plant defense (Padovan et al., 2013; Padovan et al., 2015). As yet, it is unclear whether this dramatic molecular reprogramming is the result of genetic mutations or epigenetic modifications that have accumulated amongst the mosaic branches of Eucalyptus. Most mutations are deleterious, and the long-term impact of mutational load on a slow cycling organism is more likely to lead to a “mutational meltdown” than an advantageous adaptation. Indeed, live cell imaging and surgical manipulations of axillary meristem precursors in Arabidopsis and tomato indicates that both the position and highly reduced cell division rates of these stem cell niche precursors potentially safeguard long-lived plants from mutational meltdown (Burian et al., 2016). These findings mark the tip of the iceberg in our investigation of uni-generational evolution, and raise important questions concerning the prevalence of heterogenomicity and its role in slow-cycling organismal adaptation.

## Concluding remarks

Organisms are generally assumed to function as genetically uniform individuals, but there are an overwhelming number of examples that break this assumption. Rather than being a rare phenomenon, estimations of somatic mutation rates indicate that the vast majority of multicellular organisms are by definition mosaics, and genomic uniformity is the exception rather than the norm. In an era where an organism’s genome is in essence its blueprint, the heterogenomic state forces us to re-examine how we define an individual’s genotypic identity and the relationship between genotype and phenotype. The prevalence of heterogenomicity within the plant kingdom has served both as a tool for investigating fundamental questions in developmental biology, as well as a phenomenon that generates new questions. As a tool, chimeric and mosaic analyses have demonstrated the importance of intercellular communication, context-dependent gene function, and cell lineages during tissue and organ formation. However, as a phenomenon, we are left with several open questions concerning the biological basis for heterogenomicity. We have yet to identify the intercellular signals that allow for developmental coordination across heterogenomic tissues. Furthermore, we know very little about what determines heterogenomic compatibility, and what gives rise to beneficial versus detrimental genomic combinations. Advanced molecular techniques that allow for genomic-level analyses, coupled with tissue-and cell-specific profiling, provide obvious means to begin dissecting the mechanisms underlying classic observations of communication within plants at both the organismal and intercellular levels. Chimeras and mosaics always have, and always will be, the principal method to understand intra-organismal communication during development, the cell-autonomous and non-cell autonomous activity of genes, and the consequences of heterogenomicity within an individual.

## Acknowledgements

We are grateful to Dr. Poethig for generously providing an image of the Poinsettia periclinal mosaic series (**Fig 4A**), to Dr. Elzinga and the University of California at Riverside Library for sending us a digital copy of the original ‘Bizzaria’ publication (**Fig 1A**), and to Dr. Levy for supplying us with an image of his CRISPR-Cas PSY1 genetic mosaic (**Fig 3C**). M.H.F. is supported by a postdoctoral fellowship from the NSF PGRP (award number IOS-1523669).

